# Dog Meets Man: Alloparenting and Affective Care within Co-constructed Street Dog–Human Lifeworlds in Delhi, India

**DOI:** 10.64898/2026.11.21.768014

**Authors:** Shubhangi Srivastava, Kaustabh Moni Baruah, Maan Barua, Anindya Sinha

**Author notes:** Corresponding Author: **Shubhangi Srivastava**, Animal Behaviour and Cognition Programme, National Institute of Advanced Studies, Indian Institute of Science Campus, Bengaluru 560012, India.

## Abstract

Rapid urbanisation and largely unplanned development in the cities of the Indian subcontinent have been drawing increasing ethnographic attention in recent years. Yet, even though other-than-human beings are an integral part of these spaces, the city is still most often understood from a human perspective. At the same time, nonhumans actively shape and reshape their environments through everyday behavioural choices – choices that are not only ecological, but also socioeconomic, and often influenced by their interactions and relationships with human beings. In this paper, we adopt a mixed-methods, etho-ethnographic approach to explore a curious case of other-than-human alloparenting within largely affective dog–human lifeworlds in the Indian megapolis of Delhi. We focus on a triadic relationship between two co-territorial female street dogs—*Bhuri* and *Kali*— and a particular human caregiver—*Kishan*—that reveal how these two dogs, sharing the same territory within Gur Mandi, an informal settlement in north Delhi, adjust their allo/parenting behaviour in response to their independent social interactions with the human residents of the area. This more-than-human narrative suggests that street dogs are not merely passive inhabitants of the city, but sentient, affective participants, who shape urban life through their everyday interactions. Moreover, the remarkable, counter-intuitive shifts that we observed in *multibeing* parenting behaviour point to how deeply human presence could influence and mediate short-term social interactions and the ensuing long-term relationships between nonhuman beings in their shared spaces.

## 1. PROLOGUE

It was a typical freezing winter evening for the human and canine inhabitants of a bustling market lane in the Gur Mandi informal settlement of Delhi in January 2020, just before the world spiralled into the COVID-19 pandemic (Figure 1). People, mostly men, were returning from their day jobs, the children coming out for their playtime, when several young men and women gathered around a makeshift *momo* stall. An oddly alert *Kali*^1^ – an old black-and-white female street dog – dashed across the lane and weaved her way hurriedly through a labyrinth of people to chase away a couple of adult male dogs, who had collected near the *chai/*tea shop. Barking and growling, she stared intently towards her horizon till the silhouettes of the retreating ‘foreign’ dogs disappeared against the setting sun. She quickly turned around and looked intently at a litter of seven puppies, sleeping on a rug, just around the corner of the tea shop. Panting gently, *Kali* curled around to rest on the rug, alongside the puppies, as the tea-shop owner, *Kishan* smilingly tossed some biscuits to her. Meanwhile, *Bhuri* – a young brown female street dog and the actual biological mother of the litter – quietly raised her head from a garbage dump, behind a parked motorcycle in the market lane, where she was resting and silently watched these events unfold from a distance. A curious case of other-than-human alloparenting, enmeshed within an affective dog–human lifeworld, was gently unfolding at the unlikely altar of a global, postcolonial capital city.

**Figure 1:**
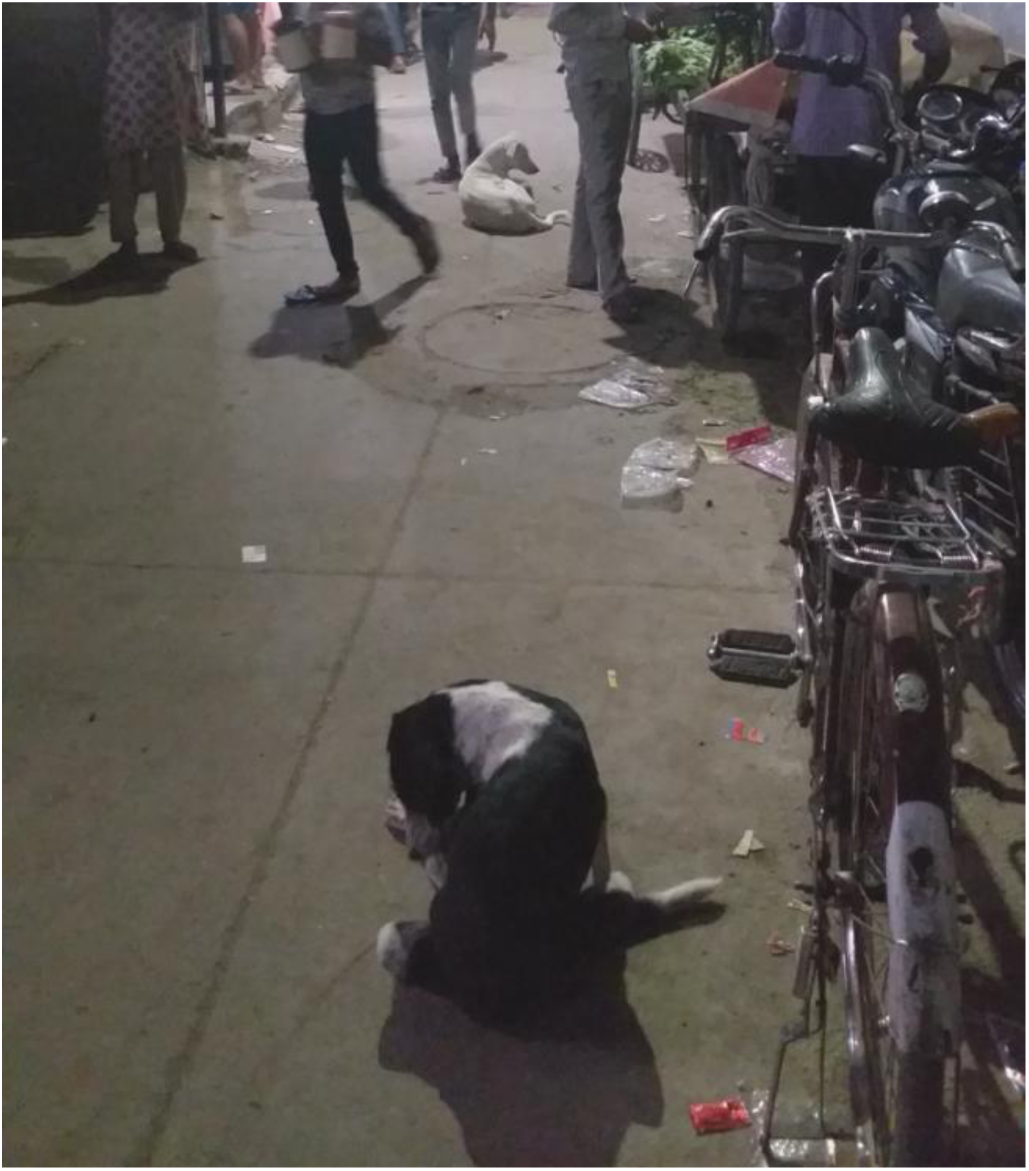
The Gur Mandi Informal Settlement in Delhi, with *Kali*, One of Our Subject Street Dogs, January 2020 **Source:** Authors’ compilation

## 2. INTRODUCTION

The rapidly growing cities of the Indian subcontinent, with their invasive urbanisation and largely unplanned development, have drawn serious ethnographic attention in recent years. The various urban processes that have been examined in Indian metropolises include the everyday struggles of the urban poor and informal settlers under the displacement plans of Delhi (Baviskar 2004, 2006); city planning, evictions, and rights to the city in Delhi (Bhan 2009), informality, urban infrastructure, and the makings of a world-class city in Kolkata (Roy 2011), public water infrastructure in Mumbai (Anand 2017), as well as mobility, gender, and city spaces in Hyderabad (Annavarapu 2021). It is noteworthy that although these ethnographic accounts of the Indian urban have often focussed on marginalised human communities, they have completely failed to explore the lives and struggles of the far more marginalised other-than-human communities and inhabitants of our cities.

Although nonhumans are integral to Indian urban spaces, the city has always been explored from the perspective of the human. With the persistent calls by certain social science and humanities scholars to reimagine cities as spaces, comparably occupied by other-than-humans globally, the almost purely anthropocentric nature of conceptualising urban spaces has been repeatedly questioned (Wolch 1996; Haraway 2008; Urbanik 2012). Urban environments have thus now been revisited as spaces, characterised by *multibeing* relationships between people, nonhumans, and other natureculture components of the city (Joshi *et al*. forthcoming; Ramakrishna *et al*. forthcoming), thus giving rise to ‘hybrid societies’, wherein humans and more-than-humans shape one another’s lives through shared histories, affects, and material entanglements within their co-constructed lifeworlds (Brunois-Pasina 2005; Lestel *et al*. 2006; Kirksey and Helmreich 2010; Barua and Sinha 2017/2019; Govindrajan 2018; Sinha and Barua 2025; Chowdhury et. al., 2024).

Such a re-conceptualisation of the city that de-centres the urban from humans as being its only inhabitants have encouraged varied descriptions of nonhuman lives in the urban. In recent years, thus, numerous studies have presented different ways of living and relating by urban nonhumans, guided, as they were, by their underlying knowledges and decision-making capacities. These studies have ranged, for example, from recognising the contributions of other-than-humans to processes of urbanisation (by chickens in Africa: Hovorka 2008; macaques in India: Barua and Sinha 2017/2019, Sinha 2025, Sinha and Barua 2025; feral cats in Canada: van Patter and Hovorka, 2018), to understanding the complexities of more-than-human relations in the urban and urbanising (Palmer 2003; Braun 2005; Power 2008; Natarajan and Sinha 2022), to examining the control of nonhuman bodies exercised by humans (Srinivasan, 2013; Narayan, 2017; Ghosh 2024) and the resulting forces of conflict, responded to and negotiated by other-than-humans (Fuentes, 2012; Knight, 2012; Barua *et al*. 2013; The Urban More-than-Human Collective 2023).

It should be noted that the surge of work on the urban nonhuman has largely been told through animal spaces—the ways in which other-than-human lives are spatially ordered, regulated, and excluded from the urban polis. Concurrently, nonhuman beings have only remained at the centre of all narratives as symbolic or material resources for humans (White and Candea 2018/2023). Nonhumans are, however, powerful agential beings and harness their multisensorial experiences to shape their own spaces in the city, in excess of human deliberations and design (Despret 2004; van Dooren *et al*. 2016; The Urban More-than-Human Collective 2023; Ragavan and Srivastava, 2020). Their dwelt environments (Ingold 2000), therefore, emerge far beyond the physical areas, designated for them by humans, to become fully fledged ‘beastly places’ that enable them to create their own belonginess deep within the city (Philo and Wilbert 2000).

Furthermore, nonhuman animals have been continually known to create and re-create their own ecologies or niches (Odling-Smee *et al*. 1996) through the socioecological decisions that they always need to make. Appropriate niche construction theories have thus been developed by behavioural ecologists to explain the effects of the immediate environment and ecological interactions on such decision-making, especially in rapidly changing urban environments (Fuentes 2010). Humans, in particular, significantly impact the lives of their fellow other-than-human beings, often implicitly, through their scientific and industrial advancements of production and consumption (Urbanik 2012). Very relevant for our current discussion, for instance, are the insights that reveal how the regular feeding of street dogs by humans could even influence and change their within-species social bonding patterns (Bhattacharjee *et al*. 2018).

In this paper, we thus provide evidence for how often-unpredictable socioecological changes in one’s immediate environment—in this case, social relations with humans— could have a profound influence on certain vital parameters of individual life-histories— here, the reproductive success of street dogs—in an intensely competitive urban situation, located in the megapolis of Delhi, in northern India. More specifically, we explore the triadic relational structure between two female dogs, *Bhuri* and *Kali*, and a human male, *Kishan*, in which the two street dogs, occupying a competitive ecological space, develop rather different relationships with *Kishan*, a local street vendor, while also maintaining vital social bonds within their pack. Street dogs are typically ‘unowned’, free-ranging, often feral individuals, found commonly across almost every street of urban India (Bhadra *et al*. 2014; CUES 2020; Barua and Sinha 2022/2023). While they are often referred to as stray dogs, they occasionally develop commensal relationships with the humans around them (Pal 2003; Bhadra *et al*. 2014). Unlike other canids like wolves or jackals, street dogs are rarely found in the ‘wild’ (but see Home *et al*. 2018) and almost always coexist with ‘their’ human communities. They, however, breed ‘naturally’, as opposed to the ‘artificially’ bred domestic dogs, also in the Indian urban (CUES 2020). This allowed us to examine, in this study based in Delhi, how the close association and social relations of certain individual street dogs with one another and with humans, all embedded within their co-constructed lifeworlds, influence and foster unusual mother–infant bonds within their community.

## 3. STUDY SITE

Delhi is a sprawling urban centre of modern India. Being the capital of a post-colonialist state, Delhi’s current nature has largely been influenced by its chequered history, especially in terms of its uneven development, governance, and the various sociocultural practices of its unique human communities. Delhi originally became the seat of political power when it was made the nation’s capital by the British colonials in 1911, while its subsequent socioeconomic and political developments followed different trajectories over the years (Shafi 2012). More specifically, Delhi’s urban development has occurred under the jurisdiction of its North, South, East, and West Municipal Corporations and New Delhi Municipal Council (DDA 2010).

Geographically, Delhi lies in the Indo-Gangetic plains, with a typically flat terrain and tropical climate. It has a human population of approximately 23 million (Jain and Mishra 2026), with about 49% of its people living in informal settlements, defined as illegal human housings lacking proper infrastructure (UN-Habitat 2015). Furthermore, Delhi has recorded an ever-growing street dog population, ranging from approximately 4,00,000 in 2016 to its estimated double of 8,00,000 in 2025 (Kumar and Sharma 2026). This burgeoning canine population has generated polarised responses of care and friction from their human co-habitants and have even formed a large part of the current political discourse in Delhi (Kumar and Sharma 2026; MCD 2026).

The actual site of our study was a busy market lane in an informal settlement – Gur Mandi (henceforth, GM) – in North Delhi (Figure 2).

**Figure 2:**
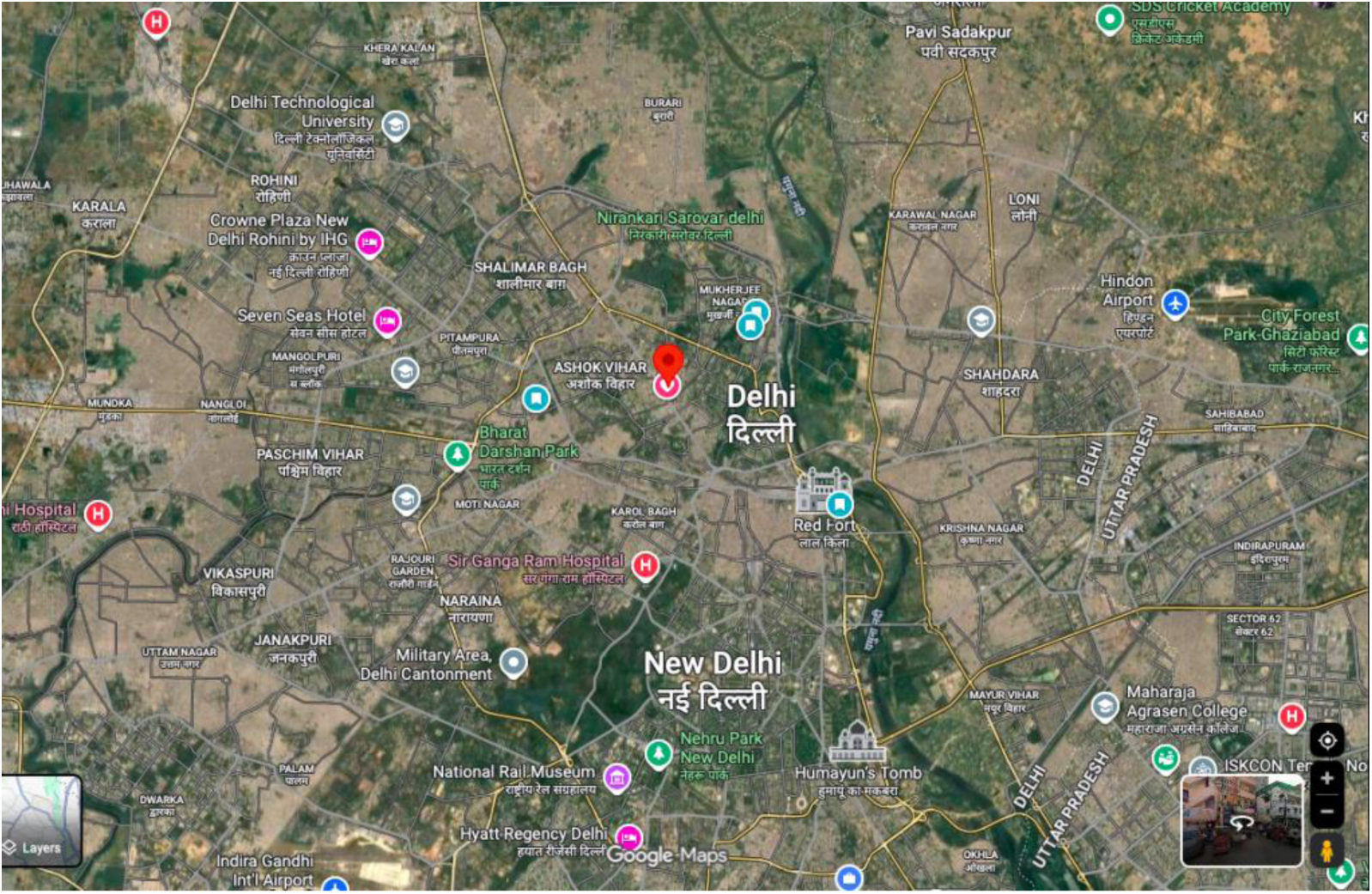
Location of Our Study Site in Delhi, January 2020 **Source:** Google Maps

GM is an urban village with a fairly populous residential area, occupied by resettled dairy farmers from the outskirts of Delhi, who were given land here by the government to rear their cattle about 50 years ago. But rapid urbanisation appeared to have compelled many of them to give up their cattle and build houses on the land, which they then rented out. The market lane, approximately 200 m long, was thus lined by several-storied houses on both sides, with food- and other shops typically on the ground floor and with residents on the floors above.

The lane was also home to 11 adult street dogs, who, according to our human respondents, have lived there for around eight years or even more, regularly interacting with the willing human residents of the street. The ambience of the area was always warm and friendly, and as soon as we reached the lane on any observation day, we would be greeted by our primary human informant, *Kishan*, originally from Uttar Pradesh, now stationed at his small tea stall, with a smiling “*Aao, chai pee lo*”—“Come, have some tea”—(Figure 3).

**Figure 3:**
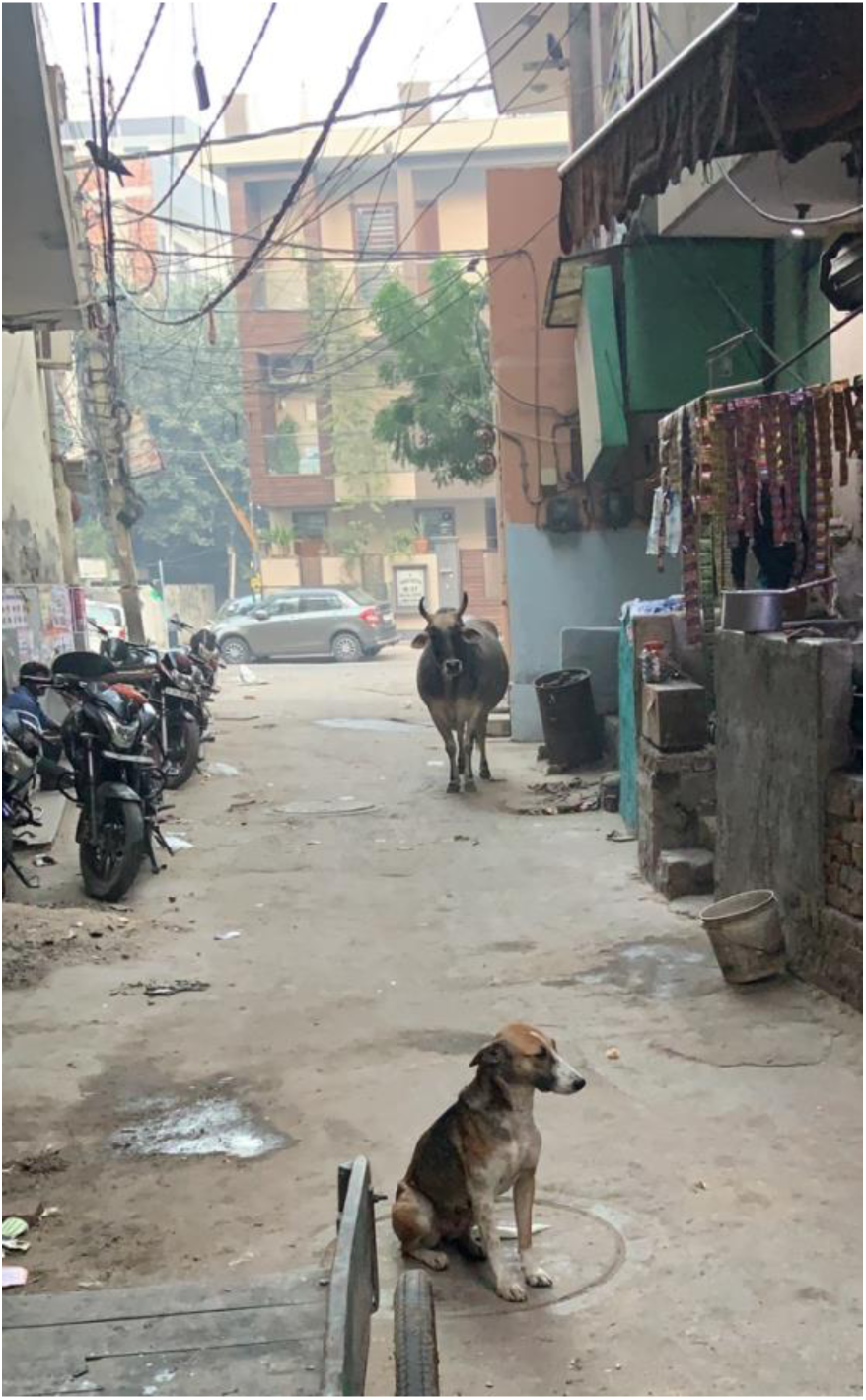
Early Morning in Gur Mandi, with *Bhuri*, One of Our Subject Street Dogs, December 2019 **Source:** Authors’ compilation

## 4. METHODS

The evolution of multispecies, multibeing studies have not been without substantial methodological challenges. By the turn of this century, both the natural and social sciences had begun to come together with their independent methodological tools to study human– nonhuman interactions and relations. Lestel and his colleagues (Lestel *et al*. 2006) pioneeringly proposed to push the boundaries of the natural and social sciences to bring forth a combined approach of etho-ethnology and ethno-ethology, which could together explore the evolutionary and cultural histories of multispecies communities. In a parallel development, innovative research on human–nonhuman relations within naturalcultural contact zones—defined as “social spaces where disparate cultures meet, clash, and grapple with each other, often in highly asymmetrical relations of domination and subordination – such as colonialism and slavery, or their aftermaths as they are lived out across the globe today” (Pratt 2008: 7) incorporated the agency of nonhuman actors and their lifeworlds in anthropology (Pratt 1991; Haraway 2008; Natarajan and Sinha 2022). From this idea of ‘becoming’ within contact zones (Haraway 2008) emerged the multispecies ethnographic approach that proposed to study the interrelated spaces and interactions of humans and other-than-humans within their co-constructed lifeworlds (van Dooren *et al*. 2016; Govindrajan 2018; Gillespie 2019). Finally, in a yet independent approach, ethnoprimatological studies have brought together primatology and ethnography to specifically study and understand human–nonhuman primate relationships (Fuentes 2010; Radhakrishna *et al*. 2012; Riley 2020). Similar to multispecies ethnography, ethnoprimatology brings forward nonhuman primates as equally situated, agential protagonists within the often-commensal spaces shared with humans, emphasising, in the process, their mutual influences and co-evolved adaptations to one another.

Even though these studies have attempted a novel, pioneering approach to comprehending multispecies lifeworlds, they have often, at worst, been considered speculative, questioned for lacking a strong foothold in ethological or behavioural observations of nonhuman lives, and critiqued for their strong claims about interspecies agency and nonhuman social decision-making not being grounded enough in the rigorous behavioural sciences (Ogden *et al*. 2013; Leth 2026). Indirectly addressing such criticisms, however, the approach of ethologically-informed ethnography (Hartigan Jr 2021) presents a methodological symmetry to studying humans and nonhumans, by systematically aligning the rigour of both systematic behavioural observations—of both human and nonhuman agents—and their ethnographic interpretations. In our paper, therefore, we have attempted to adopt a mixed-methods approach to bring together behavioural observations of street dogs in Delhi and ethnographic interpretations of their shared spaces and social interactions with humans. This particular narrative of multibeing interactions between *Bhuri, Kali,* and certain human residents, especially *Kishan*, extended from January to March 2020 and formed a part of a larger study, conducted in Delhi from May 2019 to March 2020.

The specific interactions, reported in this paper, constituted opportunistic observations within our systematic focal animal sampling—focal video recordings for 10-minutes each—on selected street dogs in urban Delhi, with *Kali* being recorded for 1 h and 34 min and *Bhuri* for 1 h and 14 min (Altmann 1974; Bonnani *et al*. 2015; Bhattacharjee *et al*. 2018). We also conducted ethnographic focal observation sessions, of one-to-two hours’ duration each, individually on *Kali* (3 h and 37 min), *Bhuri* (2 h and 40 min) and on the two of them together (1 h and 28 min), during which we richly described the behaviour and both intra- and interspecific interactions of our subject dogs (Table 1). The behavioural observations, recorded in the focal videos, were coded using an ethogram of the relevant canine behaviours (Table 2), extracted from the larger ethogram that we had earlier constructed on the basis of our observations and incorporating certain behavioural elements from earlier studies (Majumder *et al*. 2014; Pal 2005; van der Bourg *et al*. 2015).

**Table 1:** The Subject Street Dogs, Gur Mandi, Delhi, November 2019.

| **Individual** | **Name** | **Gender** | **Description** | **Group Membership** |
|---|---|---|---|---|
| 1 | *Kali* | Female | Adult, black and white pelage | BG Pack, with an adult male dog, also adopted by *Kishan* |
| 2 | *Bhuri* | Female | Adult, brown pelage | DB Pack, with three adult male dogs |

**Source:** Authors

**Table 2:** Ethogram of Canine Mothering Behaviours and Allomothering Interactions, Displayed by Street Dogs.

| **Behaviour** | **Operational Definition** |
|---|---|
| Agonism | A composite behaviour, in which several behavioural acts, including stare, snarl, bark, lunge, bite, chase, or a combination of them, are displayed by an individual towards an adversary |
| Alert | A mother or allomother displays an alert posture, with heightened attention to her surroundings, often in close association with one or more pups |
| Allogrooming | A mother or allomother grooms one or more pups with her mouth and / or her tongue |
| Block | A mother or allomother positions herself physically in the path of another individual—human or nonhuman—and prevents their movement, usually access to one or more pups |
| Nearby presence | A mother or allomother sits or moves close to one or more pups, while remaining aware of their presence |
| Pile sleep / Pile sit | A mother or allomother sleeps or sits in close bodily association with one or more pups |
| Protect | A mother or allomother displays a suite of protective behaviours, directed towards one or more pups |
| Submission | A composite behaviour, including individual behavioural acts, such as crouching or lowering of the body, tail tucking, ears flattening, avoiding of eye contact, submissive grinning, submissive urination, or a combination of them, displayed by a subordinate individual towards a dominant one |
| Suckling | The biological mother provides milk to one or more of her pups while lactating |
| Suckling attempt | A pup attempts to take their lactating biological mother's nipple in their mouth |
| Suckling rejection | A potentially lactating biological mother rejects a pup's attempt to take her nipple in their mouth |
| Watching over pups | A mother or allomother remains attentive to and watches over one or more pups |
| Withdrawal | A composite behaviour, in which an individual avoids, shrinks back, acts submissively (see above), moves away or flees from an aggressive or threatening individual |

**Source:** This study, and adapted from Pal 2005; Majumder *et al.* 2014; van der Bourg *et al.* 2015

The quantitative data on the two street dogs, which were coded using MS Excel, was analysed descriptively with respect to three behavioural situations: *Bhuri’s* nursing behaviour and its relevant chronological sequence, development and ultimate duration of maternal caregiving behaviour by *Kali*, and the directional behavioural interactions between *Kali* and *Bhuri* after the birth of the latter’s pups. Given the small sample of the opportunistically observed data, the behavioural distributions were compared using two-tailed Mann-Whitney U-tests, with sample medians, interquartile ranges, and rank-biserial correlations of the data. The focal animal sampling let our canine subjects guide us to our human respondents, with whom we conducted in-depth, open-structured interviews. *Kishan* was interviewed for a total time of about 60 min, specifically during his interactions with *Kali* and the pups, when he fed them, usually around 6:00 AM and 6:30 PM. We also conducted sporadic informal conversations with other human residents of Gur Mandi, who occasionally fed the pups. The interviews were transcribed and thematically analysed using an ethno-ethological framework, alongside the canine behavioural observations that were analysed through an etho-ethnological lens, thus considering both humans and canines as agential actors in this mutually beneficent, naturalcultural contact zone (Lestel *et al*. 2006; Natarajan and Sinha 2022). Such a mixed-methodological approach converged our simultaneously collected behavioural and ethnographic data on our human and nonhuman subjects to interpret the observed changes in canine social living, especially in response to human provisioning and other behavioural interventions.

## 5. RESULTS

Let us now return to the streets of Delhi and unravel the tripartite affective–emotional relationships that developed over the care of a set of pups between the two street dogs, *Bhuri* and *Kali*, and their human co-habitants, all occupants of the ∼300 m^2^ of a busy market lane in North Delhi (Figure 4). Of the 10 stray dogs inhabiting the area, *Bhuri*, ∼six months to one year of age, and *Kali*, more than six years old, were the only two females. *Bhuri* was a quintessential street dog, feeding from the leftovers of the neighbourhood shops while *Kali* was cared for by a tea stall owner, *Kishan*, who fed her regularly and allowed her to stay in his shop, whenever she wanted to.

**Figure 4:**
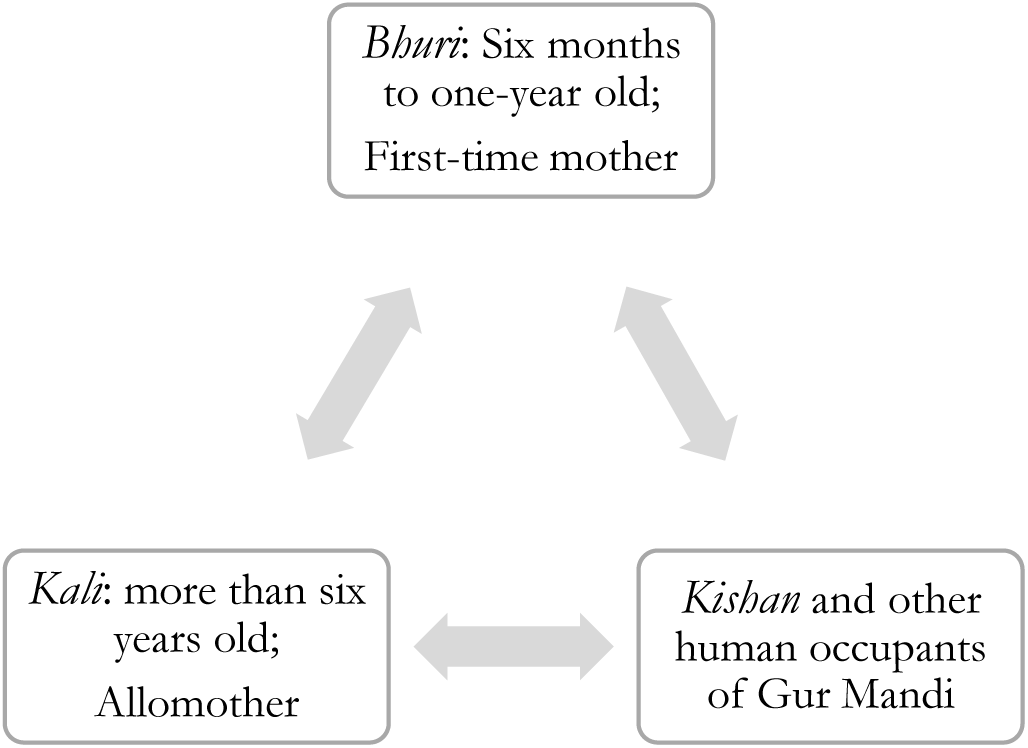
The Tripartite Relationship between the Female Street Dogs, *Bhuri* and *Kali*, and the Humans of Gur Mandi, including *Kishan*

It is also significant that *Kali* and *Bhuri* occupied independent territories prior to this study and avoided each other’s presence, when they rarely encountered one another. However, we later observed *Bhuri* to occasionally venture out to the main Gur Mandi Lane after she gave birth, as detailed below.

During the peak mating season, from October to November 2019, we observed three individual males mating with *Bhuri* during her oestrous period. *Bhuri*, who was primiparous, became a first-time mother, when she gave birth to eight pups on 13 January 2020. Until the third week of their age, the mother spent ∼70% of her time, suckling, watching over, protecting, and otherwise taking care of her pups. Subsequently, during early February, however, we observed *Bhuri* beginning to reject the suckling attempts of her pups in the nearby presence of *Kali*, occasionally even avoiding her gaze and looking away from her. This behaviour of *Bhuri*’s was inexplicable to us at that time, especially as street dogs in the Indian urban typically suckle their pups for periods ranging from six to 12 weeks (Brill *et al*. 2022).

When the pups were in their third week of age, *Kishan* started to feed them and eventually brought them over to the front of his stall, where his two children occasionally started to play with them as well. Interestingly and possibly significantly, there was a transition in the attitude of *Kali* towards the pups at this time – the first week of February – a change from being a casual, but occasionally interested, observer to becoming an active allomother, directly interacting with *Bhuri*’s pups through a range of affiliative behaviours (Figure 5). We define an alloparent as an individual, other than the genetic parent, who provides care for the offspring of other individuals, often their kin (Wilson 1975, cited by Reidman 1982). *Kali* thus began to display pile sitting or sleeping, watching over them alertly, licking and allogrooming them, and being aggressively protective of them (see Table 2 for the definitions of these behaviours). With time, over the first week of February 2020, *Kali* gradually came to dominate the physical space around the pups, tensely staring, snarling, lunging, and chasing away other dogs from the pups. More significantly, she began to display active dominance towards *Bhuri*, occasionally blocking her and preventing her access to the pups, and on occasion, threatening her with non-contact agonistic displays (Table 2).

**Figure 5:**
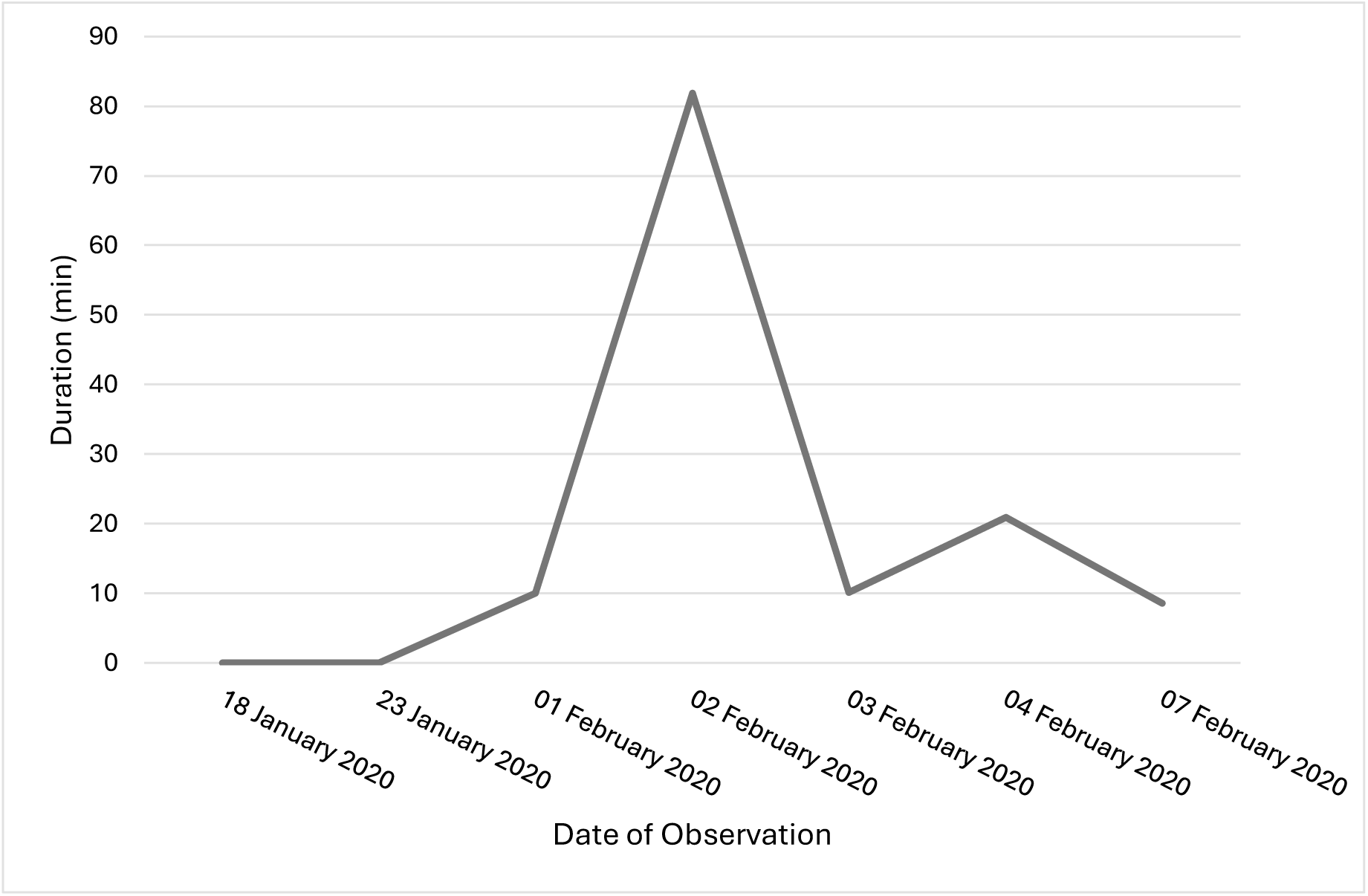
Temporal Pattern of *Kali*’s Allomothering of *Bhuri*’s Pups **Source:** Authors’ analysis **Note:** The allomothering behaviours have been described in the text.

**Figure 6:**
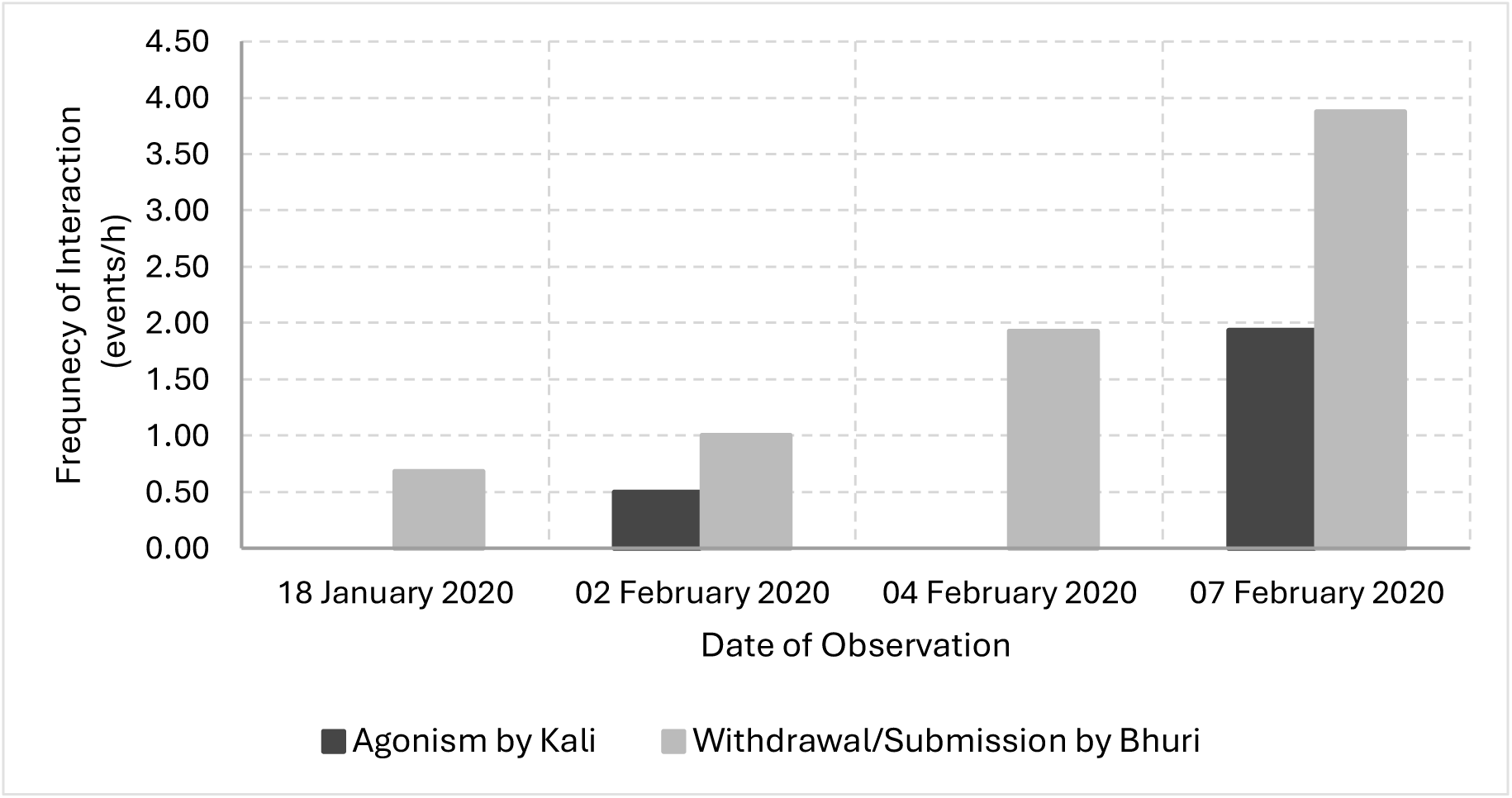
Temporal Emergence of *Kali*’s and *Bhuri*’s Agonistic–Submissive Interactions **Source:** Authors’ analysis **Note:** The agonistic and submissive behaviours have been described in Table 2.

Quantitatively, *Bhuri* spent a median of 6.30 min per 10-min session with her litter (IQR = 17.12; n = 6) while *Kali* spent a median of 10.05 min per session with *Bhuri*’s pups (IQR = 18.84; n = 6) during their respective observation periods. Although the distribution of the time spent by the two females with the pups did not differ statistically (Mann-Whitney U = 15.0, two-tailed p = 0.687; rank-biserial r = -0.17), it was clear that *Kali* spent substantial time in her allomothering of *Bhuri*’s pups. It is illuminating, however, that, as the biological mother of the pups, *Bhuri* still displayed relatively greater frequencies of certain affiliative maternal behaviours towards the pups during this time than did *Kali* (Figure 7).

**Figure 7:**
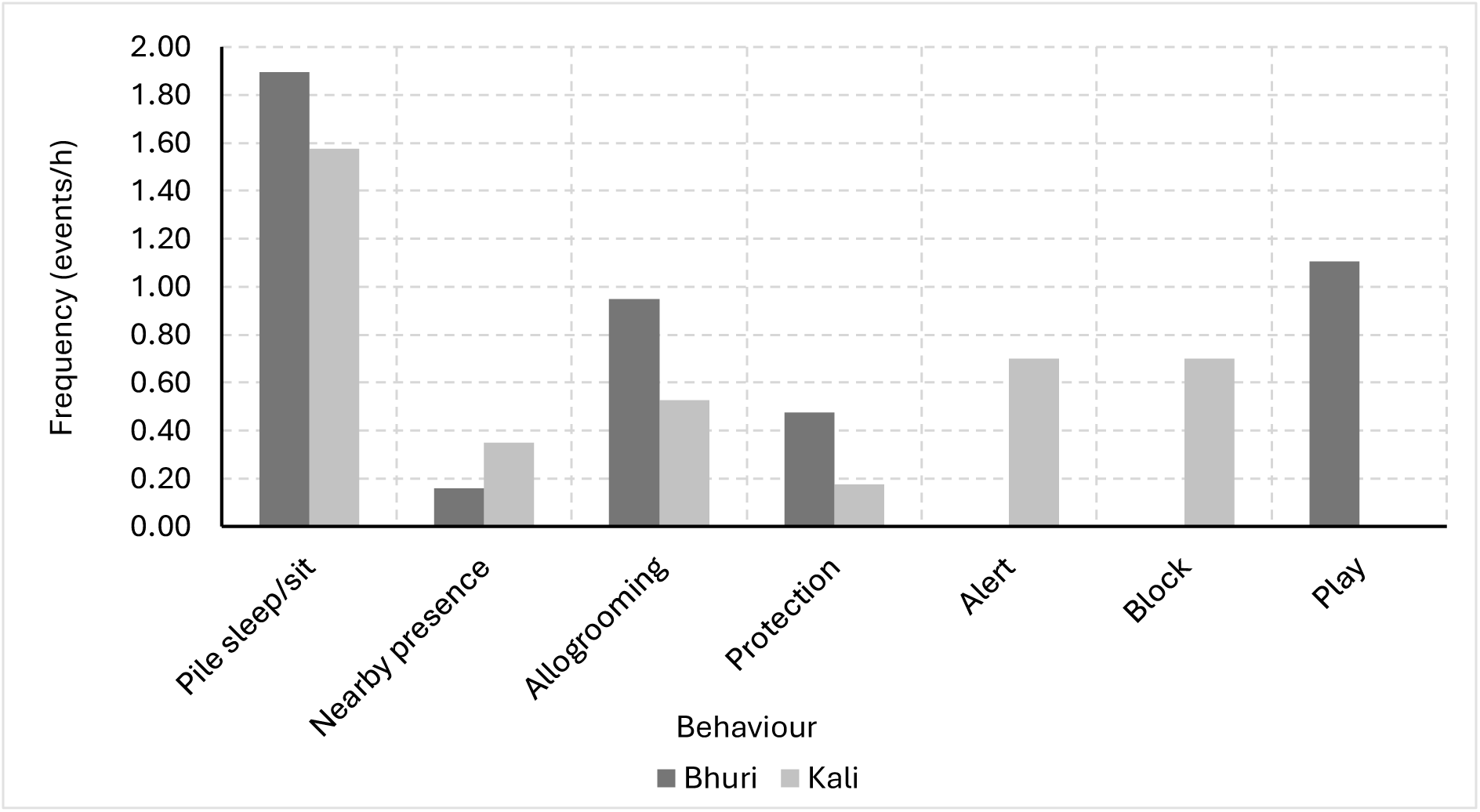
Affiliative Maternal Behaviours of *Bhuri* and *Kali* towards *Bhuri*’s Pups **Source:** Authors’ analysis **Note:** The affiliative behaviours have been described in Table 2.

**Figure 7:**
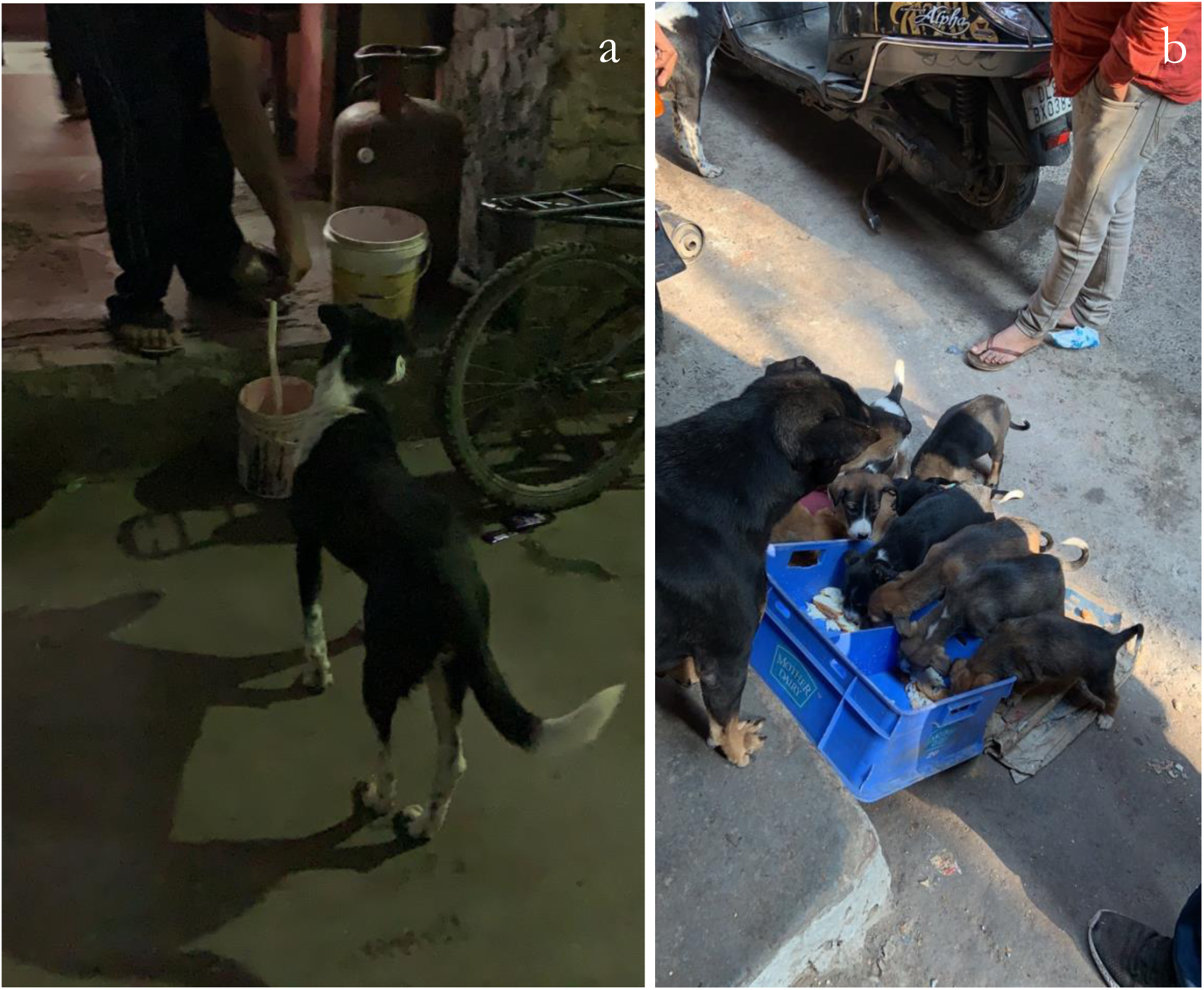
Interspecies Affiliation: (a) *Kisan* Provisioning *Kali*, and (b) Human Residents of Gur Mandi Feeding *Bhuri*’s Pups while *Kali* Looks on Alertly **Source:** Authors’ compilation

A third participant of this multibeing encounter were the human occupants of the area, some of whose everyday interventions in the quotidian lives of other-than-human beings could alters their social structures, behavioural patterns or their life-history strategies. As a case in point, *Kali* and *Bhuri* were subjected to strikingly differential treatment by the local people. While *Kali* was generally adored and hence, often artificially provisioned, *Bhuri* was relegated to indifference, thus largely dependent on human-source garbage for food. This was highlighted in a discussion with *Kishan*, wherein he opined that *Bhuri* did not feed her pups adequately and claimed that this possibly made *Kali* a more appropriate mother to *Bhuri*’s pups. A very different picture, possibly significant from the perspective of gender, emerged when, on one occasion, we observed Pushpa, a middle-aged woman resident, driving *Kali* away from the pups so that *Bhuri* could comfortably return to nursing her litter. There is now increasing evidence, emerging from integrated multi-methods approaches like ours, that such intricately nebulous configurations within human–nonhuman lifeworlds, as exemplified by the tripartite relationships between *Bhuri*, *Kali*, and the human residents of Gur Mandi, could result in novel multispecies behavioural developments—in this case, the unique juxtaposition of Kali’s rapidly altering antagonistic and protective behaviours—especially in highly dynamic and anthropogenic settings (Barua and Sinha 2022/2023).

In this connection, we speculate that the unusual allomothering behaviour of *Kali* could have manifested successfully, partly in response to the strong affective and emotional stimuli, generated by the unique attachment that simultaneously developed between *Kali* and *Kishan*, on the one hand (Figure 8) and between *Bhuri*’s pups and their human caregivers, including *Kishan*, his two children, and other humans in the area (Figure 9). This, we believe, could have influenced and led *Kali* to care for *Bhuri*’s pups as her ‘own’ *kin*. These stimuli could have been further strengthened by *Kishan’s* belief, formed from his own lived experience and imagined lifeworlds, about *Kali* being the ‘true’ mother of the pups. While talking about *Bhuri*, the biological mother of the pups, for example, he once said, as indicated earlier, “*My Kali is a better mother than Bhuri*. *She does not even feed the pups*”. In contrast, Pushpa, who had chased away *Kali* from the pups, opined, “*The black one does not ever let the actual mother come near the pups because she had lost her pups once and hasn’t been able to give birth since. I always kick her away so that the pups can be near their own mother*.”

## 6. POSTSCRIPT

We saw later, in April 2022, that only a single individual male, from *Bhuri*’s litter of eight pups, had survived to maturity. He was named *Titu* and taken care of by *Kishan* and his family, along with *Kali*.

*Bhuri* and *Kali* had also come to occupy their usual territories in different parts of Gur Mandi, where we had seen them prior to this study, at that time.

## 7. DISCUSSION

### 7.1 Alloparental Care

Alloparenting, typically defined as parental care that is provided by individuals towards the offspring of other individuals, often genetically related to them, is a universal behaviour amongst humans. It is believed that this behaviour may have considerably influenced our evolutionary history and continues to remain important even in contemporary society (Kenkel *et al*. 2017). Interestingly, alloparenting is primarily limited to only 3% of nonhuman mammalian species (Russel 2004; Kenkel *et al*. 2017). Reidman (1982) reports that alloparenting is mostly observed in comparatively large-bodied K-selected species, which maintain a stable population size in predictable ecosystems, with intense competition for limited resources, focusing, in the process, on high parental investment in a relatively few offspring that they produce over their lifetime but ensuring a high probability of their survival. Many species of the genus *Canis*, especially wolves, coyotes, wild dogs, and domestic dogs have now been established to practice extended parental care, often with the help of alloparents (Mazumdar *et al*. 2014).

It is noteworthy, however, that alloparental care appears to be rarely seen in free-ranging street dogs, as compared to wild and domestic dogs. It has been hypothesised that this absence could be attributed to evolutionary changes in the foraging ecology of free-ranging dogs from being active hunters to more settled scavengers, which, in turn, may have resulted in heightened intra-group competition over food resources and reduced cooperation amongst them while domestic dogs may exhibit more pronounced cooperative tendencies, including alloparenting, owing to the more regular presence of stable food sources (Pongracz and Sztruhala 2019). This argument has been furthered by Bhadra *et al*. (2015), who observed the presence of high-quality food substrates in the environment increasing the frequency of parent–offspring conflict in free-ranging dogs, as mothers attempt to ensure higher lifetime reproductive success rate by even competing with their current litter for food (Bhadra *et al*. 2015). Despite the infrequent alloparenting, shown by free-ranging dog populations, however, some instances of alloparenting by grandmothers have occasionally been reported in Indian street dogs (Pal *et al*. 2014).

While existing studies on alloparental care in street dogs suggest that the phenomenon is restricted to kin in efforts to decrease infant mortality and simultaneously increase inclusive fitness (Mazumdar *et al*. 2014), our etho-ethnological observations challenge this view. We thus report an apparently evolutionary incongruity: a female street dog, *Kali* alloparenting a litter of possibly non-kin pups, born to *Bhuri*, defying the established biological drive of kin selection. *Kali* was socially dominant to *Bhuri* and extensively displayed agonism towards her, especially when competitively taking care of the pups. The complete absence of any affiliative interactions between the two individuals during our long-term study, in contrast, suggests an absence of any cooperative relationship between the two dogs and thus argues against the phenomena of mutualism: direct, immediate mutual benefits to cooperating individuals, or reciprocal altruism: delayed return of benefits to the participating individuals, as ultimate causes of *Kali*’s allomothering (Trivers 1971; Axelrod and Hamilton 1981; Boucher 1985; West *et al*. 2007).

Several canid species, such as gray wolves, dingoes, African wild dogs, and free-ranging street dogs, are cooperative breeders that often raise altricial litters in dens. Although infrequent, agonistic competition, involving non-kin allomothering, manifests in these species primarily around coerced allomothering, allo-nursing, den access or forced redirection of care. In coerced allomothering, for example, a dominant female could eliminate a subordinate female’s litter and then coerce the subordinate female into allo-nursing and provide care for the dominant individual’s surviving pups (African wild dogs: Frame *et al*. 1979; dingoes: Corbett 1988; Indian street dogs: Pal 2005; gray wolves: Lukas and Huchard 2019). Again, in some cases, including in Indian street dogs, unrelated low-ranking females could perform pup-guarding and provisioning as an appeasement strategy to avoid violent eviction by the dominant female. Here, alloparental behaviour is maintained by the constant threat of agonistic aggression (Malcolm and Marten 1982; Sillero-Zubiri *et al*. 1996; Macdonald *et al*. 2004; Pal *et al*. 2021). In all these cases, however, it is the dominant female that benefits by forcing allomothering of her pups by subordinate females in a variety of socioecological situations.

Conversely, the unusual appropriation of a subordinate, non-kin female’s litter by a non-lactating dominant female, coupled with the observed active exclusion of the biological mother—which we observed in this study—could represent, we hypothesise, a form of non-lethal reproductive interference, functionally analogous to “aunting-to-death”, reported in nonhuman primates (Hrdy 1976). In resource-limited urban environments, monopolising a rival’s pups, without providing milk, we believe, could induce neonatal starvation and eliminate future resource competitors without the energetic costs or injury risks of direct infanticidal combat, unless buffered by anthropogenic provisioning (Paul et al. 2014, 2016; Paul and Bhadra 2018). Proximately, this behaviour may have been driven by lingering maternal neurochemistry, following an earlier litter loss—possibly the situation with *Kali*, as reported by Pushpa—or by covert pseudopregnancy, wherein elevated prolactin and oxytocin could maintain compulsive brooding motivation despite the cessation or absence of lactation (Lezama-García *et al*. 2019; Santos *et al*. 2020; Gobello 2021). As central oxytocin simultaneously amplifies defensive aggression against perceived intruders, this may have driven *Kali* to aggressively guard the appropriated pups while repeatedly driving off *Bhuri* (Bosch and Neumann 2012).

An alternative cognitive hypothesis would suggest that spatial-temporal kin-recognition rules could have also caused *Kali* to misattribute pups, held within her core territory, as her own, and this may have been further promoted by her social bonding with *Kishan*, as noted earlier. Simultaneously, *Kali* may have lacked the causal foresight to recognise that non-lactating guarding of pups could inevitably result in high litter mortality, as also happened with *Bhuri*’s allomothered pups (Holmes and Sherman 1982; Miklósi 2015). Thus, proximate hormonal priming, cognitive constraints, and an active perception and internalisation of a strong interspecific bond could have, interestingly, produced a synergetic, though agonistic, outcome, functionally equivalent to infanticide.

### 7.2 Affective Exchanges

Moving away from an examination of alloparenting from a unidimensional ethological perspective, we now explore the fabric of entanglement between human and other-than-human beings and attempt to understand the affective exchanges underlying and shaping human–nonhuman lifeworlds in highly anthropogenic urban habitats.

Deeply entangled human–nonhuman relationships, especially those unfolding within co-constructed hybrid lifeworlds, have often been scrutinised from the perspective of affect (Sinha *et al*., 2021; Banerjee and Sinha 2023; Ramakrishna *et al*. 2023), following the *affective turn* in human and nonhuman geography (Thien 2005; Koivunen 2010; Pile 2010). More-than-human studies, seeking to engage seriously with other-than-human lives, has historically turned to the natural sciences. Here, we deepen our etho-geographical dialogue by investigating the analytical value that social science concepts – most notably, affect – could bring to our ethological interpretations. Defined as an ‘intensity’ (Massumi 2002), affect constitutes a sentient singularity, unbound from conventional systems of representation. Circulating within and across both human and other-than-human bodies, affects draw beings together along shared pathways and durations. Simultaneously enacted and experienced, they can also be generated, choreographed or deliberately engineered. As much quotidian as they are extraordinary, affects shape the cadences of everyday life and the patterned routines of dwelling (Stewart 2007). Consequently, affect differs fundamentally from emotion: while emotions, alongside cognitivist framings of affect, remain objects of the conscious mind, affect operates on a non- or pre-cognitive register and not generally expressed or understood through language (Ahmed 2010; Pile 2010). It is this understanding of affect—as a transpersonal (Anderson 2006) and trans-species capacity (Despret 2004)—that anchors our current analysis.

Within more-than-human geographies and affective ethologies, this pre-cognitive, trans-species capacity is further articulated as the dynamic, embodied ability of both human and nonhuman bodies to affect and be affected through situated interspecies encounters (Sinha *et al*. 2021). Rather than requiring direct access to internal animal subjectivity, an affective lens attends to the shifting relational processes, sensory exchanges, and behavioural dispositions through which sentient beings interpret, respond to, and co-compose shared lifeworlds (Sinha *et al*. 2021). In contexts of ecological entanglement, therefore, affect operates as an experiential mode of ‘reading’ and engaging with nonhuman lives: shaping how humans and animals attune to one another’s agency, bodily presence, and ecological histories (Banerjee and Sinha 2023). Furthermore, these fluid affective states intersect with sociocultural narratives and emotional responses, generating shifting ‘affectual geographies’ that govern tolerance, negotiation, and coexistence across contested human– nonhuman interfaces (Sinha *et al*. 2021; Ramakrishna et al. 2023). It is also noteworthy that recent multispecies ethnographies have begun to focus, in rich detail, on actual human–nonhuman short-term encounters and long-term relations, the abilities of individual nonhumans to shape such interactions and relationships, and most importantly, to foreground the affective agency and subjectivities of the other-than-human beings themselves (Bear 2011; Watson and Warkentin 2013; see Srinivasan 2013, 2019 and Narayanan 2017 for agency in free-living/street dogs), thus highlighting the nature of *multibeing*—rather than simply multispecies—relationships (Ramakrishna *et al*. forthcoming).

In developing nations of the Global South, like India, streets abound in a variety of local canine breeds. *Bhuri* and *Kali* are, in particular, Indian pariah dogs, a landrace of free-ranging street dogs, native to the Indian subcontinent, with a characteristic wedge-shaped head, erect ears, and a curved tail. This breed is an example of an ancient group of dogs, commonly known as *pye-dogs* (from *pāhī* in Hindi, translating to ‘outsider’) that live in or close to human settlements throughout South Asia, South China, the Middle East and North Africa (Vessey-Fitzgerald 1957/2023). There is now archaeological evidence that the Indian street dog was present in ancient Indian villages even as early as 4,500 years ago (Sewell and Guha 1931; Prashad 1936).

More relevant for our discussion, virtually all Indians are aware of the extent and nature of the relationships that Indian street dogs have been involved in with urban citizens across the country over generations and the space of their own they have created in city streets through their interactions with humans in their everyday lives. These free-ranging, commensal dogs have increasingly become dependent for their survival on humans not only for provisioned food resources, but also for their wellbeing on their relationships with humans (Bhadra 2014; Majumder *et al*. 2014; Bhattacharjee *et al*. 2017; Nandi *et al*., 2025). The availability and type of food resources (the regular provisioning of *Kali* and of *Bhuri*’s pups by *Kishan*, this study) and the strength and quality of the affiliative interspecies relationships that develop between individual dogs and certain humans *in situ* (“*meri Kali*” or “my *Kali*,” *Kishan*, this study) could thus profoundly impact the social structure, prevailing mating and pup-rearing patterns, and inter-individual sociability within local street dog communities (Pal 2003; Bhattacharjee and Bhadra 2020).

When applied to the street dogs of urban India, our affective framework illuminates how the survival outcomes and reproductive dynamics of individual dogs are co-constituted through embodied encounters between particular dogs, certain human residents, and shared street spaces (Sinha *et al*. 2021; Barua and Sinha 2022/2023). In our particular observed case of litter appropriation, the non-lactating dominant female *Kali*’s takeover of a subordinate female *Bhuri*’s pups and her repeated exclusion of the biological mother illustrate how intense, embodied affective forces—potentially mediated by hormonal priming and neonatal sensory cues—can drive non-cooperative allomothering, even at the cost of litter mortality. Both dogs, therefore, appear to have actively ‘read’ and navigated the prevailing human affective dispositions towards them, thus attuning their denning choices and foraging routines to local gestures of tolerance or hostility (Banerjee and Sinha 2023). Additionally, *Kali* was perhaps able to comprehend and ultimately depend on the strong affective—and potentially emotional—familial bonds, expressed by *Kishan* and his children towards her; we also believe that this long-term, inter-generational, interspecies social relationship may have been read by *Kali*, when manifest in the form of their daily affective exchanges of care. Furthermore, we argue that *Kali* was motivated to take over *Bhuri*’s litter at an opportune moment, when the pups were introduced into her territory by *Kishan* and his affective care extended to them. *Kali*, a member of a highly territorial species, may have then decided to include the pups under her care as an extension of her social bonding with *Kishan*, re-awakening her maternal instincts soon after losing her own pups. *Kishan*’s affective support for *Kali*’s new-found responsibility may have, in turn, led him to describe her to as “the actual mother [of the pups],” who “performs motherly duties better [than *Bhuri*]” to us. Was *Kali* also able to read this affective belief of *Kishan*’s through their daily exchanges and could this transmitted belief encouraged her allomothering activities?

Finally, it is possible that *Kishan*’s provisioning and caregiving of both *Kali* and the pups may have relieved *Kali* of the urgent need to lactate and otherwise provision the pups, thus buffering, at least to a little extent—the ultimate survival of a single pup, *Titu*—the lethal fitness consequences of the non-lactating *Kali*’s allomothering of the pups. Thus, the behavioural trajectories and reproductive fates of our subject Indian street dogs emerge from these fluid, multibeing, affectual geographies, wherein canine agonism and human care continually intersect, with occasionally critically functional consequences.

## 8. CONCLUSIONS

Street dogs in the Indian capital city of Delhi have increasingly become subjects of public anxiety and political debate, with human feeders and caregivers frequently bearing the brunt of calls for the complete removal of dogs from the Indian urban (Saha 2025). Yet, sustained human provisioning, often motivated by deeply held beliefs of receiving *punya*, ‘a diffuse, cleansing merit from God’ (Sinha *et al*. 2021), generates vital food-distribution networks and ‘affective corridors’ that guide human–canine movement as soft infrastructures, running counter to formal urban planning (Kumar 2026). Extending this perspective, our study demonstrates how these *multibeing* affective relations shape dogs not merely as social actors, but as biological beings. Through an interdisciplinary etho-ethnographic methodology that examines shared streets as natural laboratories (Bhadra 2014), we have documented a hybrid human–dog society, in which regular human provisioning altered innate alloparental behaviour. Specifically, the tripartite relationship amongst *Kali*, *Bhuri*, and *Kishan* revealed to us a very unusual case of non-kin alloparenting, marked by agonism between the biological mother, *Bhuri* and the alloparent, *Kali*. Relieved of foraging pressures by regular feeding and socially bonded to her long-term benefactor, *Kishan* through sustained affective exchanges, *Kali* extended her care towards *Bhuri*’s pups despite the energetic costs, illustrating how interspecies affective and emotional entanglements can reshape fundamental canid behavioural ecology.

Viewing the city through these emergent affective relations shifts urban analysis from concrete structures of roads and buildings to lively infrastructures of co-constructed lifeworlds of multispecies exchanges (Simone 2004; Barua and Sinha 2017/2019, 2022/2023). Recognising street dogs as sentient co-creators of the urban fabric thus expands more-than-human urbanism and affirms the city as a *zoöpolis* (Wolch 1996). Over the long term, our findings, we believe, carry critical implications for both more-than-human lives in the city and urban governance. Given the extent to which human care has traditionally and purposefully modified the social and reproductive behaviours of street dogs across the Indian urban, contemporary conflicts cannot be resolved simply through exclusionary removal of street dogs. Instead, sustainable coexistence requires governing and supporting street-level networks of affective care alongside humane population management strategies, such as Animal Birth Control (ABC) programmes. Ultimately, acknowledging how deeply human provisioning and emotional bonds have altered canine biology and sociality in India over generations is essential for the negotiated forging of ethically grounded, resilient multispecies cities in the disruptive Anthropocene of today.

## ACKNOWLEDGEMENTS

The idea for this manuscript began after a field interview with *Kishan*, our principal human respondent and we, therefore, express our sincere gratitude to him for his time and endless cups of tea. We also owe a huge debt of gratitude to the nonhuman participants of this study, whose presence and interactions made this inquiry possible but without ever seeking their consent. SS would like to thank Shruti Ragavan for her valuable comments on an earlier draft of our manuscript.

## Funding

This study was funded by a European Research Council Horizon 2020 Starting Grant, entitled *Urban Ecologies: Governing Nonhuman Life in Global Cities* (Grant Number 759239). It also forms part of the doctoral thesis of SS under the same project grant.

## Author Contributions

SS, AS, and MB conceptualised and designed the study; SS and KSB collected primary field data; SS and AS analysed the data; SS wrote the first draft of the manuscript, with contributions from KSB; SS and AS edited and revised the manuscript; MB and AS supervised the entire study. All authors read and approved the final manuscript.

## Ethics Statement

We hereby confirm that this study complies with all requirements of ethical approvals from the Research Ethics Committee of the National Institute of Advanced Studies in Bangalore, India for the conduct of this research.

## Data Availability Statement

The data used to support the findings of this study are available from the corresponding author upon reasonable request.

## Conflict of Interest Statement

The authors declare no competing interests, financial or otherwise, in the conduct of this study.

## Footnotes

1 The names of human respondents have been changed for use in this study. However, the names of the canine respondents are same as those given by the human residents.

## Notes

### Competing Interest Statement

The authors have declared no competing interest.

